# Hepatocyte-, but not myeloid cell- Rictor/mTORC2 deficiency moderately attenuates steatotic liver disease induced by intake of a choline-deficient, amino acid-defined high-fat diet

**DOI:** 10.64898/2026.06.17.732842

**Authors:** Bianca F. Leonardi, Ana B. Pires, Marina A. Abe-Honda, Loreana Silveira, Álbert S. Peixoto, Érique Castro, Thayna S. Vieira, Natália M. Pessoa, Erika V. M. Pessoa, Natália Pontara-Corte, Guo Yin, Marlene S. Kohlhepp, Ana Carolina P. Baptista, Mariana Mesquita, Helayne S. de Freitas, Camila N. Bezerra, Frank Tacke, Adrien Guillot, William T. Festuccia

## Abstract

Previous studies have demonstrated that mechanistic target of rapamycin complex 2 (mTORC2) deficiency provides complete protection against steatotic liver disease driven by constitutive activation of the phosphoinositide 3-kinase (PI3K)–Akt signaling pathway and de novo lipogenesis, and partial protection against disease induced by a high-fat diet. We investigated herein whether mTORC2 deficiency in hepatocytes and myeloid cells, including Kupffer cells and recruited macrophages, influences the development of liver disease induced by intake of a choline-deficient, amino acid-defined high-fat diet (CDAHFD), a model in which liver disease is induced by impaired hepatic secretion of very low-density lipoprotein (VLDL) triacylglycerol. For this, mice with either hepatocyte- or myeloid cells-specific deletion of mTORC2 essential component rapamycin-insensitive companion of mTOR (Rictor) and their respective littermate controls were fed with either chow or CDAHFD for 10 weeks and evaluated for hepatic steatosis, inflammation and fibrosis. Our main findings indicate that hepatocyte Rictor/mTORC2 deficiency slightly attenuated the CDAHFD-induced increases in liver mass, macrovesicular steatosis and triacylglycerol accumulation, without affecting though liver cholesterol, serum markers of liver injury (AST and ALT), as well as the upregulation in proinflammatory cytokine IL-1β and expression of fibrosis-related genes. Myeloid cells-Rictor deletion had no detectable impact on liver steatosis, inflammatory, or fibrosis induced by CDAHFD. In conclusion, mTORC2 deficiency show modest beneficial effects in counteracting liver disease induced by CDAHFD intake.

## Introduction

Metabolic dysfunction-associated steatotic liver disease (MASLD) encompasses a wide spectrum of highly prevalent hepatic diseases with varying degrees of severity, ranging from a relatively harmless steatosis to the more severe metabolic dysfunction-associated steatohepatitis (MASH), fibrosis, cirrhosis, and ultimately MASH-related hepatocellular carcinoma (HCC)^1^. MASLD global prevalence is estimated at approximately 30%, a number that is expected to increase in the next decades, underscoring the urgent need for effective strategies to prevent disease development and progression^2^. Although MASLD is currently the most prevalent form of chronic liver disease^3^, only two drugs, the thyroid hormone receptor β ligand resmetirom and the glucagon like peptide 1 (GLP-1) analog semaglutide, have recently been approved for MASH treatment^4,5^. Thus, a better understanding of the molecular mechanisms underlying MASH pathogenesis could help in the identification of novel therapeutic targets to expand the spectrum of strategies to prevent disease development and progression.

MASLD pathophysiology involves a complex interplay between genetic, environmental, and metabolic factors that altogether promote hepatic steatosis, lipotoxicity, oxidative stress, and insulin resistance, major disease hallmarks that interconnectedly drive the development of liver inflammation, injury, and fibrosis^6^. MASLD lipid accumulation and steatosis occur when rates of hepatic *de novo* lipogenesis (DNL) and free fatty acid influx from adipose tissue lipolysis, both of which markedly increased in patients with MASLD/MASH due to insulin resistance, exceeds rates of fatty acid oxidation and secretion as very low-density lipoprotein (VLDL)-triacylglycerol^7^.

Previous studies have demonstrated that the mechanistic target of rapamycin complex 2 (mTORC2) plays a critical role in the regulation of liver DNL and MASLD development. Indeed, mTORC2 inactivation through deletion of its essential component rapamycin independent companion of mTOR (Rictor) in hepatocytes completely and partially protected mice from liver disease induced by constitutive activation of hepatocytes phosphoinositide 3-kinase (PI3K)-Akt signaling and intake of a high fat diet, respectively^8^. Indeed, hepatocyte mTORC2 deficiency reduced liver sterol regulatory element-binding protein-1c (SREBP1c) transcriptional activity, glycolysis, and DNL, leading to a decrease in liver mass, in part due to a reduction in hepatic glycogen and triacylglycerol contents^8^. mTORC2 is one of the two multiprotein complexes that share the serine/threonine kinase mechanistic target of rapamycin (mTOR) as catalytic core. Primarily activated by growth factors such as insulin, mTORC2 coordinates cell survival, metabolism, and cytoskeletal organization through the regulation of several members of the AGC family of kinases, including Akt, serum- and glucocorticoid-induced kinase 1 (SGK1), and protein kinase C-α (PKC-α)^9^. Beyond its critical role in hepatic DNL regulation, mTORC2 was shown to be activated by toll-like receptor (TLR) ligands and functions as a key negative regulator of intracellular proinflammatory signaling in macrophage^10^. Notably, hepatic macrophages, including resident Kupffer cells and recruited monocytes, are central drivers of MASLD progression, orchestrating lipid-induced inflammation, hepatocellular injury, and fibrogenesis^11^. Despite of mTORC2 prominent role in MASLD induced by enhanced DNL, as well as counteracting macrophage proinflammatory responses, little is known about whether mTORC2 signaling in hepatocytes and macrophages influences MASLD development, particularly in murine models in which disease progression is not driven by DNL.

In this context, the choline-deficient, L-amino acid-defined, high-fat diet (CDAHFD), containing 60 kcal% fat and low methionine (0.1% by weight), is a well-established model of progressive liver steatosis and fibrosis. Choline deficiency impairs VLDL synthesis and secretion, promoting intrahepatic triacylglycerol accumulation, while the high dietary fat content further aggravates lipotoxicity and metabolic stress, thereby accelerating liver injury and fibrogenesis^12^. Therefore, considering the pivotal role of hepatocytes and macrophages to MASLD pathogenesis and the involvement of mTORC2 in metabolic and inflammatory regulation, this study aimed to define the cell-specific contribution of mTORC2 to the development of CDAHFD-induced MASLD.

## Methods

### Mice

Experimental procedures involving animals were approved by the Animal Care Committee of the Institute of Biomedical Sciences, University of Sao Paulo, Brazil (#115/2016 and 6160250820, CEUA). All mice were on a C57BL/6J background and obtained from Jackson Laboratories (USA). Hepatocyte-specific Rictor deletion was generated by crossing Rictor Lox/Lox (rictor^tm1.1Klg/SjmJ^) mice with albumin-cre mice (B6.Cg-*Speer6-ps1Tg(Alb-cre)21Mgn*/J). Myeloid cells (macrophages, neutrophils, etc)-specific Rictor deletion was generated by crossing Rictor Lox/Lox (rictor^tm1.1Klg/SjmJ^) mice with LyzM-cre mice (B6.129P2-Lyz2tm1(cre)If). Male Rictor Lox/Lox; Albumin-Cre^+^ mice (hereafter referred to as L-RicKO) and their littermate Rictor Lox/Lox controls (L-RicWT) were used for the hepatocyte-specific experiments. Male Rictor Lox/Lox; Lysozyme M-Cre^+^ mice (hereafter referred to as M-RicKO) and their littermate Rictor Lox/Lox controls (M-RicWT) were used for the macrophage-specific experiments. Mice were maintained at 23 ± 1°C, 12:12 h light-dark cycle, with free access to water and food ad libitum. From 8 weeks of age, mice were fed either a nonpurified chow diet (70% carbohydrate, 20% protein, 10% fat, in %kcal, NUVILAB CR-1®-Sogorb Inc., Paraná, Brazil) or a high-fat, choline-deficient amino acid-defined diet (CDAHFD, 20% carbohydrate, 20% protein, 60% fat in % kcal, PragSoluções) for 10 weeks. Body composition was evaluated using nuclear magnetic resonance spectroscopy (Bruker Minispec). Mice were anesthetized with isoflurane and euthanized by cervical dislocation for tissue and blood collection. Genotypes were determined by PCR analysis of tail genomic DNA.

### Liver histology

Liver samples were fixed in 4% paraformaldehyde and embedded in paraffin. Sections 5.0 μm-thick were obtained and stained with hematoxylin and eosin (H/E) to assess general liver morphology.

### Serum determinations

Serum glucose, triacylglycerol, total cholesterol, alanine aminotransferase (ALT) and aspartate aminotransferase (AST) activities were measured using commercially available kits according to the manufacturers’ instructions (Labtest, Lagoa Santa, Brazil).

### Liver composition

Liver samples were homogenized in buffer containing 140 mM NaCl, 50 mM Tris (pH 7.4), 0,1% Triton X-100, resuspended in isopropanol, and evaluated for triacylglycerol and cholesterol content with colorimetric kits (Labtest, Lagoa Santa, Brazil). Hepatic glycogen content was determined by alkaline hydrolysis as previously described^13^. Briefly, liver (50 mg) was digested in 30% KOH saturated with Na_2_SO_4_ and glycogen was precipitated by centrifugation after addition of ethanol. Pellet was washed, dissolved in water, and hydrolyzed with 300 µL of 4N H_2_SO_4_ at 70°C. Samples were subsequently cooled and neutralized with 300 µL of 4 N NaOH. Glucose concentration was determined using a colorimetric kit (Labtest, Lagoa Santa, Brazil).

### Acetate oxidation and incorporation in TAG-fatty acids (DNL) in liver slices

Liver slices of 1000 μm produced with a McIlwain Tissue Chopper were incubated in hermetically-closed vials, containing 1.5 mL of Krebs-Ringer Bicarbonate buffer pH 7.4 composed of (in mM) 5 glucose, 0.51 MgCl_2_, 4.56 KCl, 119.8 NaCl, 0.7 Na_2_HPO_4_, 1.3 NaH_2_PO_4_, 15.0 NaHCO_3_, 2% essentially fatty acid-free albumin, containing 2 mM acetate and 0.2 μCi/vial [1-^14^C] acetate for evaluation of fatty acid synthesis (*de novo* lipogenesis) and carbon flux in the tricarboxylic acid (TCA) cycle. After 2 h at 37°C, the medium was acidified and labeled CO_2_ was trapped in phenylethylamine-ethanol-moistened piece of paper and liver slices were washed with saline and destined for lipid extraction with chloroform/methanol for measurement of acetate incorporation into triacylglycerol-fatty acids, as described^14,15^.

### Citrate Synthase Activity

Liver was homogenized in 50 mM phosphate buffer and centrifuged at 13,000 rpm for 10 min. The supernatant was incubated with 0.1 M Tris–HCl, 500 μM oxaloacetate, 200 μM 5,50-dithiobis(2-nitrobenzoic acid) (DTNB) and 100 μM acetyl-CoA for 5 min at 30°C. DTNB reduction was followed at 412 nm as previously described^16^.

### Hepatic cytokine content

Liver samples (100 mg) were homogenized in 500 µL of extraction buffer. Protein concentration in the supernatant was determined by the Bradford method^17^ using bovine serum albumin as standard (Bio-Rad, Hercules, CA, USA). Hepatic concentrations of tumor necrosis factor alpha (TNF-α), interleukins IL-1β, IL-6, and IL-10 were quantified by ELISA (DuoSet ELISA, R&D Systems, Minneapolis, MN, USA) according to the manufacturer’s instructions.

### RNA extraction and qPCR

Total RNA was extracted from liver samples (50 mg), reverse transcribed into cDNA, and subjected to quantitative PCR analysis as previously described^18^. Primer nucleotide sequences are depicted in Table 1. Relative gene expression was calculated using the 2^-ΔΔCt method. Data were normalized to the housekeeping gene cyclophilin, whose expression was not significantly affected by genotype or experimental conditions.

**Table 1.**
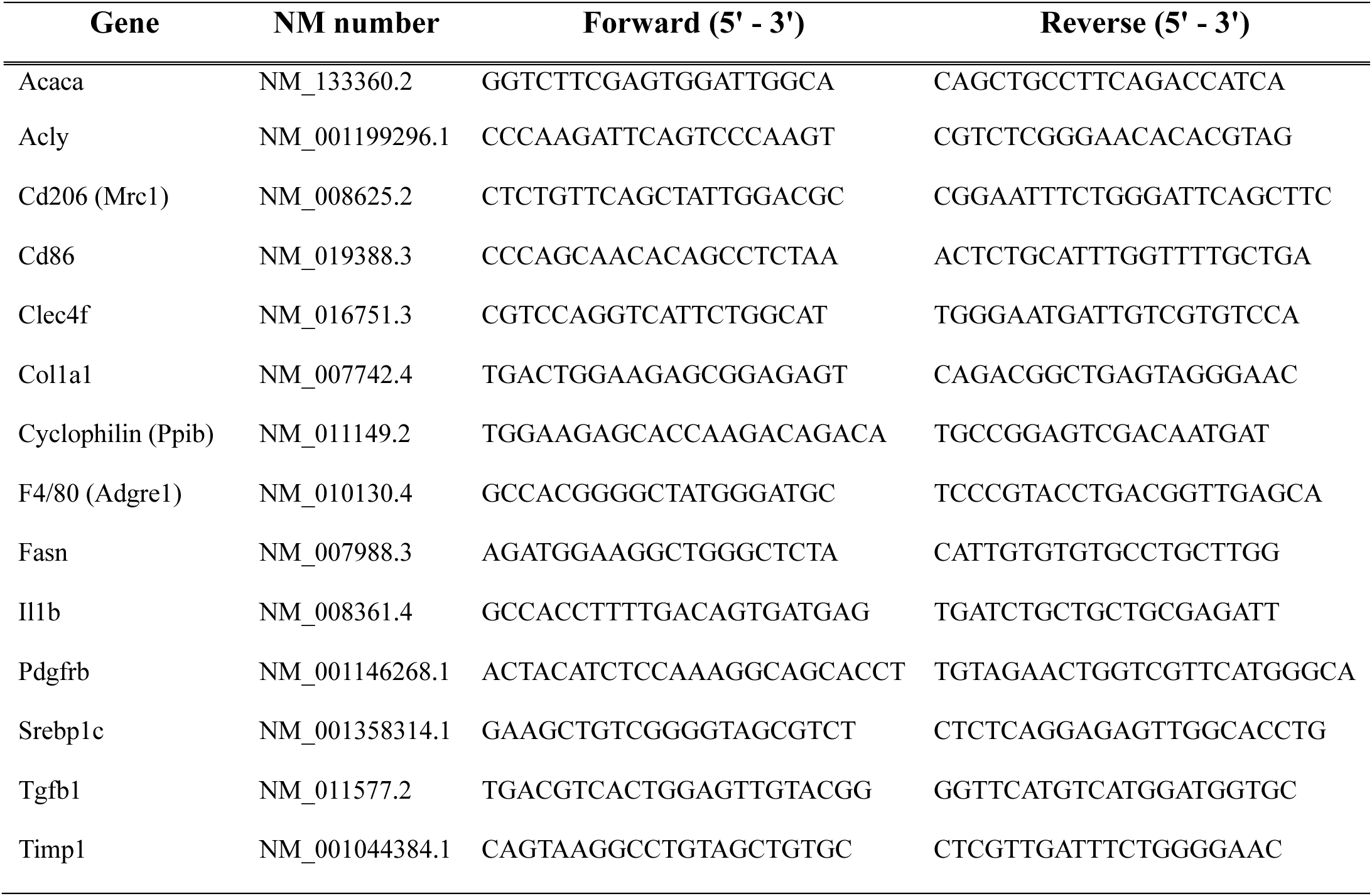
Primers used in qPCR.

### Western Blotting

Liver samples were homogenized in buffer containing 50 mM HEPES, 2 mM EDTA, 10 mM sodium pyrophosphate, 40 mM NaCl, 10 mM sodium glycerophosphate, 50 mM NaF, 2 mM sodium orthovanadate, 1% Triton-X 100, and EDTA-free protease inhibitors, centrifuged, and analyzed for protein content as previously described^18^. Primary antibody against mitochondrial oxidative phosphorylation (OXPHOS) complexes (Abcam ab110413) was diluted at 1:1000 in TBS-T 5% milk. Membranes were incubated with peroxidase-conjugated secondary antibody (1:5000) and revealed with ECL-enhanced chemiluminescence substrate (GE Healthcare).

### Multiplex Immunohistochemistry

Sequential multiplex immunofluorescence was performed on 2 μm thick Formalin-Fixed Paraffin-Embedded (FFPE) mice liver sections as previously described^19,20,21^. The antibody elution buffer was prepared from distilled water, 0.5 M Tris–HCl (pH 6.8), 10% (w/v) sodium dodecyl sulfate, and 2-mercaptoethanol. To remove artifacts due to autofluorescence from diet components, the BG555 signal was subtracted from IBA1 and CLEC4F images obtained from CDAHFD samples prior to analysis.

### Statistical analysis

Data are presented as mean ± SEM. Statistical analyses were performed using two-way ANOVA followed by Tukey post hoc test for multiple comparisons. Differences were considered statistically significant when p < 0.05. Analyses were conducted using GraphPad Prism (GraphPad Software, San Diego, CA, USA).

## Results

As illustrated in Figure 1A-C, intake of a CDAHFD for 10 weeks promoted body weight loss and reduced lean mass without significantly impacting fat mass in comparison to chow-fed mice, such effects that were independent of mice genotype. Extending previous findings^8^, chow-fed L-RicKO mice displayed reduced liver mass when compared to littermate L-RicWT on the same diet. Intake of CDAHFD increased liver mass in both L-RicWT and KO mice, such an effect that was of higher magnitude in the former than the latter (Fig. 1D). Along with liver mass, CDAHFD intake promoted liver macrovesicular steatosis in both L-RicWT and KO mice as evidenced by the enhanced ballooning found in H/E staining of liver sections and the increased hepatic contents of triacylglycerol and cholesterol (Fig. 1E-G). Noteworthy, the upregulation in liver ballooning and triacylglycerol content induced by CDAHFD was of higher magnitude in L-RicWT than KO mice, while the increase in cholesterol content was similar in both genotypes (Fig. 1E-G). In addition to lipids, both hepatocyte Rictor/mTORC2 deficiency and CDAHFD intake independently reduced liver glycogen and protein contents, while their combination did not further affect glycogen, but additively decreased liver protein content (Fig. 1H and I). CDAHFD intake significantly increased serum levels of liver injury markers aspartate and alanine amino aminotransferases (AST and ALT), independently of mice genotype (Fig. 1J-K). Serum glucose and cholesterol levels were reduced by CDAHFD in L-RicWT, but not L-RicKO mice, when compared to chow-fed mice, whereas serum triacylglycerol was not altered by either CDAHFD intake, mice genotype or their combination (Fig. 1L–N).

**Figure 1.**
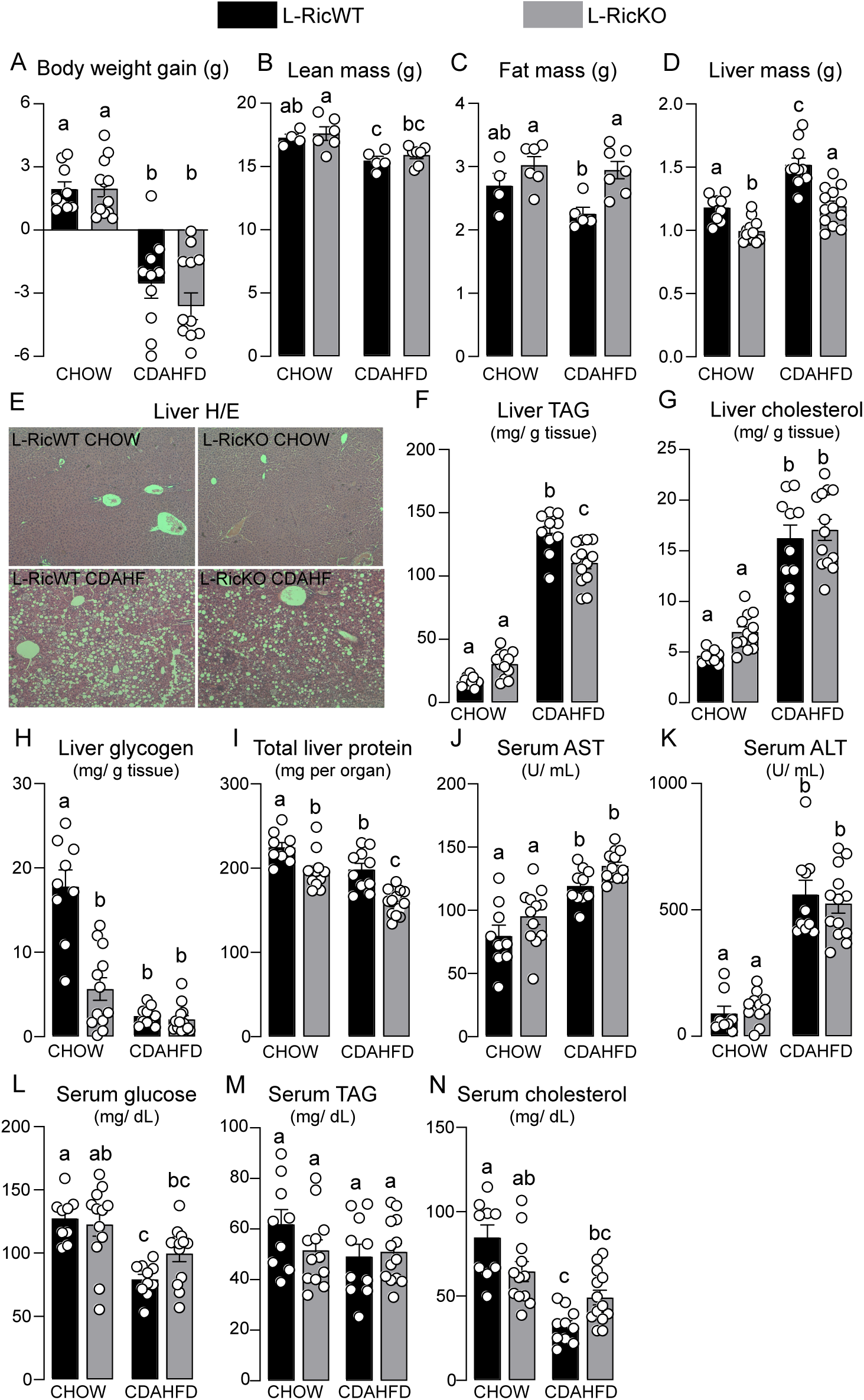
Loss of mTORC2 in hepatocytes reduces liver mass and hepatic triacylglycerol (TAG), but not glycogen or total cholesterol (TC) content, in mice fed a CDAHFD for 10 weeks. Body weight gain (A), lean mass (B), fat mass (C), liver mass (D), representative H&E-stained liver sections (E), hepatic triacylglycerol (TAG, F), total cholesterol (TC, G), glycogen (H), and protein (I) contents, as well as serum aspartate aminotransferase (AST, J), alanine aminotransferase (ALT, K), glucose (L), TAG (M), and TC (N) levels in male Rictor flox (L-RicWT) and Rictor flox albumin Cre^+^ (L-RicKO) mice fed either chow or choline-deficient, L-amino acid-defined, high-fat diet (CDAHFD) for 10 weeks. Data are presented as mean ± SEM. Two-way ANOVA followed by Tukey’s multiple-comparisons test. Groups that do not share a common superscript letter are significantly different (p < 0.05).

Next, we investigated the mechanisms underlying the reduced hepatic lipid accumulation observed in L-RicKO mice fed with CDAHFD. As illustrated in Figure 2A, both hepatocyte Rictor/mTORC2 deficiency and CDAHFD intake independently reduced hepatic DNL evaluated through the incorporation of ^14^C-acetate into triacylglycerol in liver slices, without additive effects of their combination. Similarly to DNL, both hepatocyte Rictor/mTORC2 deficiency and CDAHFD intake independently reduced mRNA levels of key enzymes/proteins involved in DNL namely *Acly* and *Srebf1*, while CDAHFD intake, but not hepatocyte Rictor/mTORC2 deficiency significantly reduced liver mRNA content of *Acaca* and *Fasn* (Fig. 2B-E). There were no additive effects of hepatocyte Rictor/mTORC2 deficiency and CDAHFD intake on the mRNA content of these proteins (Fig. 2B-E). We also assessed liver oxidative metabolism by measuring carbon flux in the TCA cycle with labeled acetate, along with citrate synthase activity, a surrogate of mitochondrial mass, and protein content of mitochondrial oxidative phosphorylation (OXPHOS) complexes. CDAHFD intake significantly increased acetate oxidation in L-RicKO to levels that were similar to those found in CDAHFD-fed L-RicWT (Fig. 2F). Furthermore, CDAHFD-fed L-RicWT mice featured higher citrate synthase activity than CDAHFD-fed L-RicKO mice (Fig. 2G). No effects of genotype, diet and their combination were observed in the contents of complexes I and III, while chow-fed L-RicKO mice featured a higher content of complex II when compared to chow-fed L-RicWT, such a difference that was fully abolished upon CDAHFD intake (Fig. 2H-J). Finally, CDAHFD intake significantly reduced liver complex V regardless of mice genotype (Fig. 2K).

**Figure 2.**
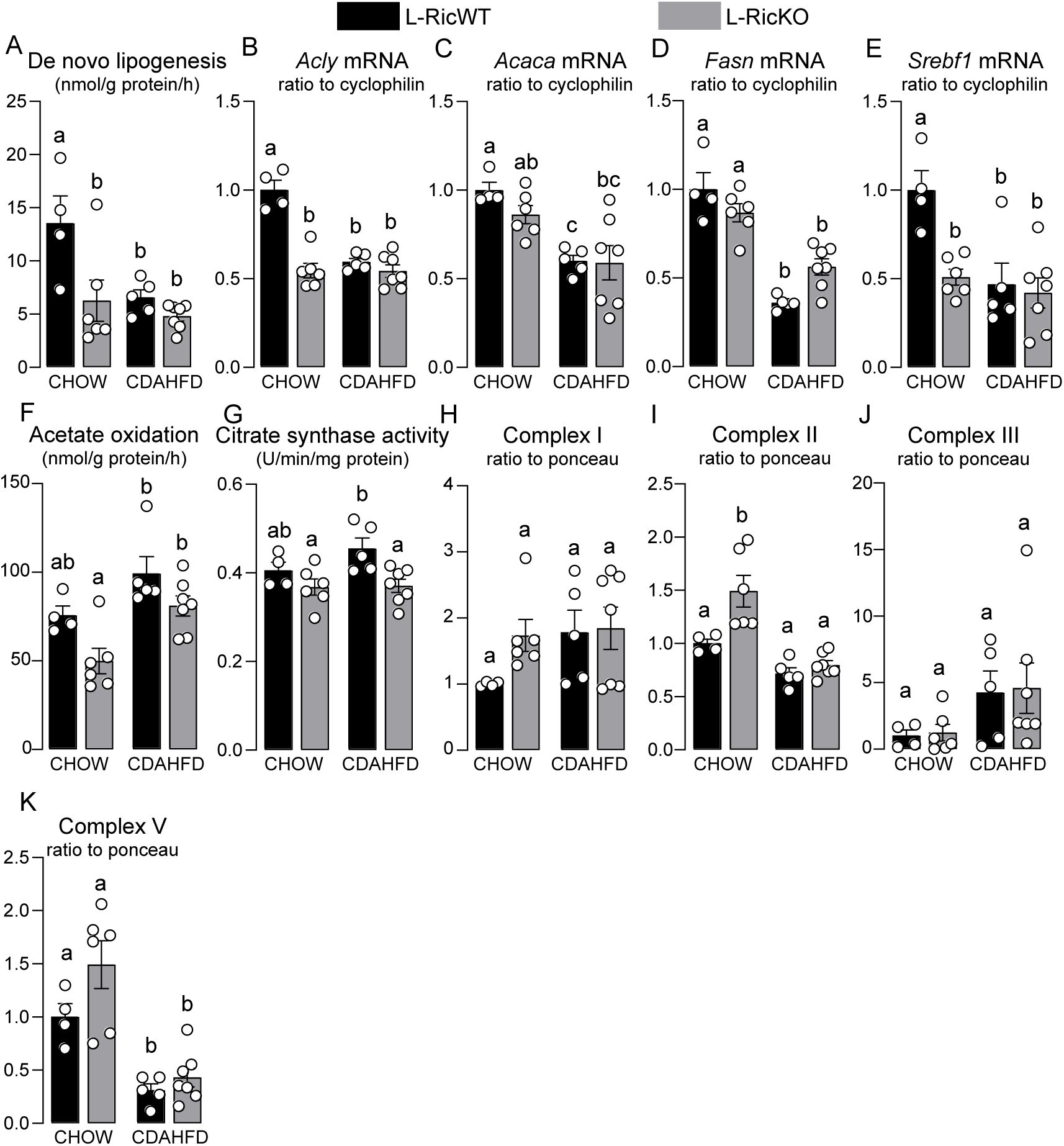
CDAHFD intake and hepatocyte mTORC2 deficiency reduce hepatic *de novo* lipogenesis (DNL) independently, without additive effects. Incorporation of [^14^C]-acetate into triacylglycerol-fatty acids (A); hepatic gene expression of the DNL-related enzymes ATP citrate lyase (*Acly*, B), acetyl CoA carboxylase (*Acaca*, C), fatty acid synthase (*Fasn*, D), and sterol regulatory element binding factors 1 (*Srebf1*, E); hepatic oxidation of [^14^C]-acetate (F); citrate synthase activity (G); and mitochondrial respiratory complex I (H), II (I), III (J), and V (K) contents in male Rictor flox (L-RicWT) and Rictor flox albumin Cre^+^ (L-RicKO) mice fed either chow or choline-deficient, L-amino acid-defined, high-fat diet (CDAHFD) for 10 weeks. Data are presented as mean ± SEM. Two-way ANOVA followed by Tukey’s multiple-comparisons test. Groups that do not share a common superscript letter are significantly different (p < 0.05).

Inflammation is a key driver of MASH development and progression toward more severe stages of liver disease, which are characterized by disruption of hepatic architecture and collagen deposition resulting from hepatic stellate cell activation. Indeed, CDAHFD intake significantly increased hepatic IL-1β mRNA and protein content in both L-RicWT and KO mice (Fig. 3A–B). In contrast, CDAHFD intake reduced liver content of the proinflammatory cytokines TNF-α and IL-6 and the anti-inflammatory cytokine IL-10 independently of mice genotype (Fig. 3C–E). Along with *IL-1β*, CDAHFD feeding increased liver mRNA content of *Adgre1* (*F4/80*), a general marker of mature macrophages, and *Cd86*, a marker of pro-inflammatory (M1) macrophages, regardless of mice genotype. In contrast, liver mRNA content of *Clec4f*, a marker of Kupffer cells, was unaltered by diet, genotype and their combination, while CDAHFD feeding increased the liver mRNA content of *Mrc1*, a marker of alternatively activated (M2) macrophages in L-RicKO mice to levels that were similar to those found in CDAHFD-fed L-RicWT (Fig. 3F–I). Furthermore, liver content of fibrosis-related genes, including *Col1a1*, *Tgfb1*, *Pdgfrb*, and *Timp1*, was markedly increased in CDAHFD-fed mice, regardless of genotype (Fig. 3J–M).

**Figure 3.**
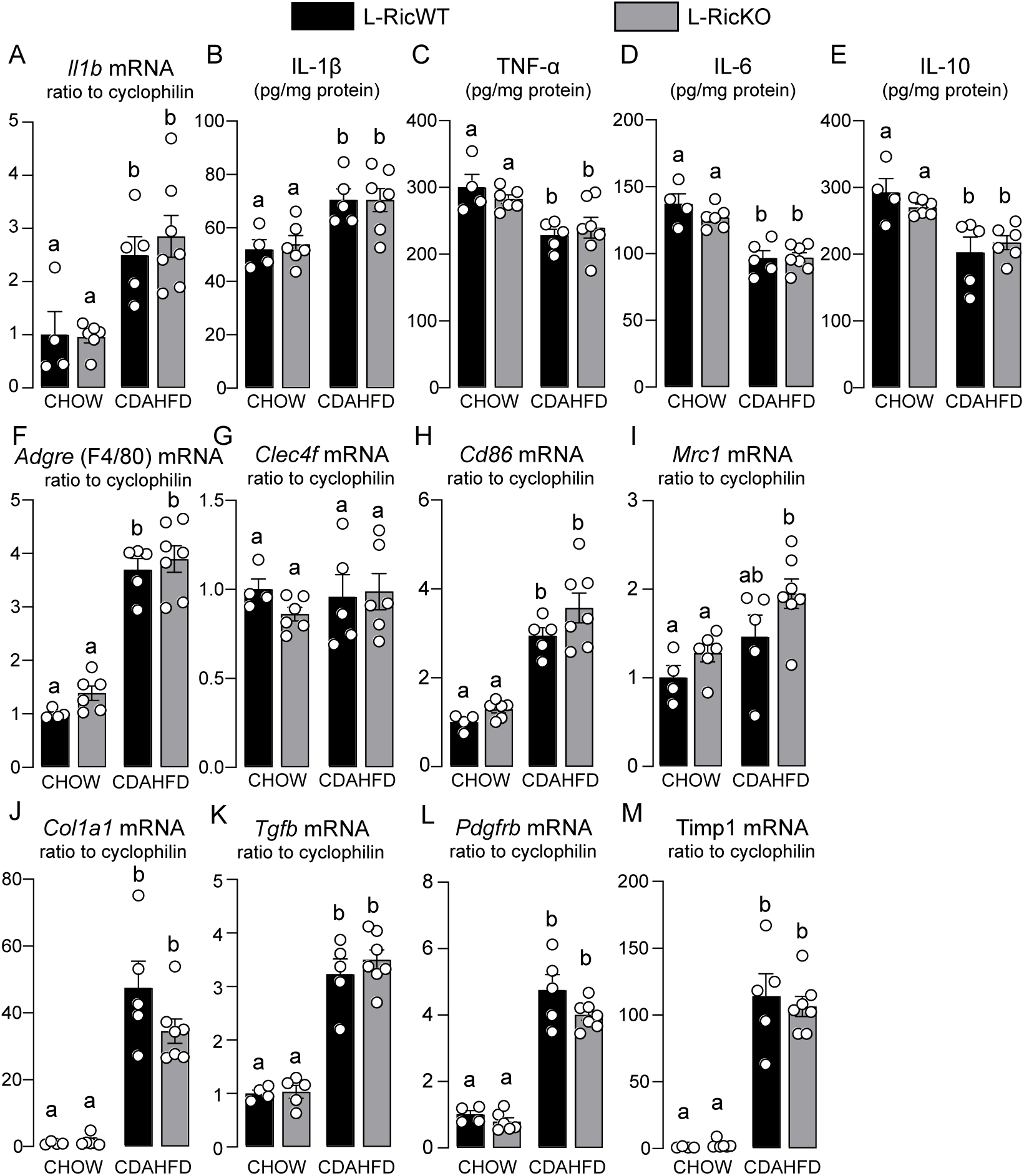
CDAHFD intake increases liver macrophage accumulation, IL-1β, pro-fibrotic gene expression regardless of mice genotype. Hepatic interleukin 1β (*Il1b*) gene expression (A) and protein content (B), tumor necrosis factor α (TNFα, C), IL-6 (D) and IL-10 (E) protein content and hepatic gene expression of *Adgre* F4/80 (F), *Clec4f* (G), *Cd86* (H), mannose Receptor C-Type 1 (*Mcr1*, I), collagen 1a1 (*Col1a1*, J), transforming growth factor β (*Tgfb*, K), *platelet-derived growth factor receptor* β (*Pdgfrb*, L) and tissue inhibitor of metalloproteinases (*Timp1*, M) in male Rictor flox (L-RicWT) and Rictor flox albumin Cre^+^ (L-RicKO) mice fed either chow or choline-deficient, L-amino acid-defined, high-fat diet (CDAHFD) for 10 weeks. Data are presented as mean ± SEM. Two-way ANOVA followed by Tukey’s multiple-comparisons test. Groups that do not share a common superscript letter are significantly different (p < 0.05).

To gain deeper insight into the spatial distribution of hepatic cell populations, particularly liver macrophages, we performed multiplex immunofluorescence staining. L-RicKO mice displayed a higher density of DAPI-positive nuclei than L-RicWT upon chow intake. Furthermore, CDAHFD feeding increased cellular density in both genotypes, as evidenced by the higher number of DAPI-positive cells per unit area compared with their respective chow-fed counterparts (Fig. 4A). Regarding macrophage populations, the total number of IBA1^+^ cells, a pan-macrophage marker, did not differ significantly among the experimental groups (Fig. 4D). CDAHFD intake, however, was associated in L-RicWT mice with a trend toward increased liver abundance of IBA1⁺CLEC4F⁻ cells and a significant increase in IBA1⁺TIMD4⁻ cells, two populations indicative of recruited macrophages, in comparison to chow-fed L-RicWT. In contrast, L-RicKO mice fed with CDAHFD exhibited a tendency toward reduced numbers of both IBA1⁺CLEC4F⁻ cells (L-RicWT CDAHFD vs. L-RicKO CDAHFD; p = 0.1087) and IBA1⁺TIMD4⁻ cells (L-RicWT CDAHFD vs. L-RicKO CDAHFD; p = 0.0837), suggesting a modest attenuation of macrophage recruitment in the absence of Rictor (Fig. 4B-C). In contrast, the IBA1⁺CLEC4F⁺ and IBA1⁺TIMD4⁺ populations, representing Kupffer cell-like macrophages, were reduced in CDAHFD-fed mice regardless of genotype (Fig. 4E-F;H-I). PCNA, a marker of cellular proliferation, was similarly increased in CDAHFD-fed mice across both genotypes (Fig. 4G). In addition, CDAHFD intake increased cholangiocytes CK7^+^ cells in L-RicWT, but not L-RicKO mice (Fig. 4GJ. Co-staining for PCNA and either IBA1 or CK7 revealed no evidence of increased macrophage proliferation (PCNA^+^IBA1^+^ cells). In contrast, CDAHFD intake increased proliferation of biliary epithelial cells (PCNA^+^CK7^+^ cells), an effect that was observed exclusively in L-RicWT mice fed (Fig. 4K-L).

**Figure 4.**
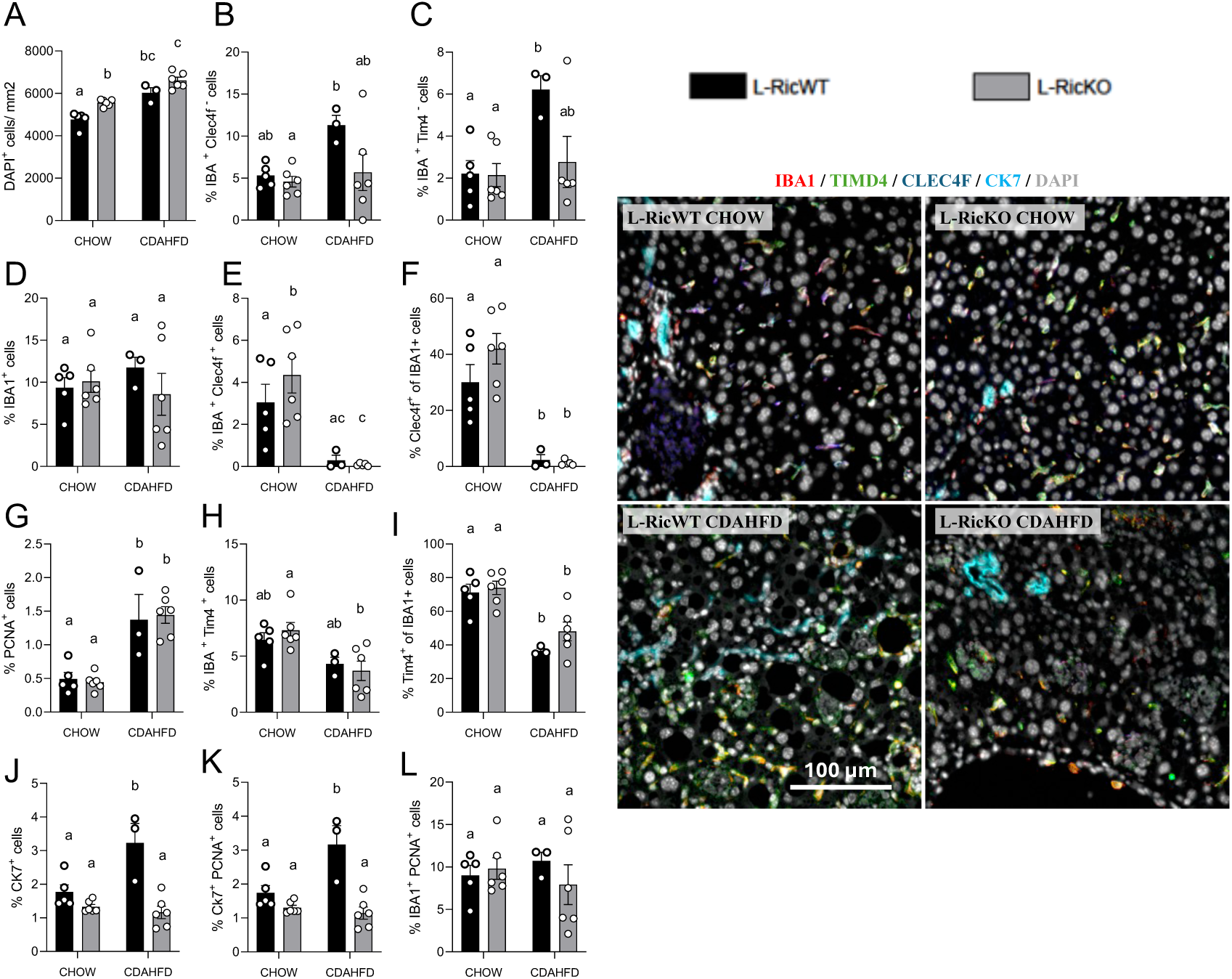
Multiplex immunofluorescence reveals that CDAHFD intake increases hepatic macrophage infiltration and decreases Kupffer cells, with minor effects of mTORC2 deficiency. Immunofluorescence staining for Ionized calcium-binding adaptor molecule 1 (IBA1), T cell immunoglobulin and mucin domain containing 4 (TIMD4), C-Type Lectin Domain Family 4 Member F (CLEC4F), cytokeratin 7 (CK7) and proliferating cell nuclear antigen (PCNA) in liver sections of male Rictor flox (L-RicWT) and Rictor flox albumin Cre^+^ (L-RicKO) mice fed either chow or choline-deficient, L-amino acid-defined, high-fat diet (CDAHFD) for 10 weeks. DAPI-positive cells per mm2 of tissue (A); % of total cells that are IBA1-positiveCLEC4F-negative (B), IBA1-positiveTIMD4-negative (C), IBA1-positive (D), IBA1-positiveCLEC4F-positive (E), PCNA-positive (G), IBA1-positiveTIMD4-positive (H), CK7-positive (J), CK7-positivePCNA-positive (K), IBA1-positivePCNA-positive (L); % of IBA1-positive cells that are CLEC4F-positive (F) or TIMD4-positive (I). IBA1 and CLEC4F staining in CDAHFD samples, the BG555 background signal was subtracted. Data are presented as mean ± SEM. Two-way ANOVA followed by Tukey’s multiple-comparisons test. Groups that do not share a common superscript letter are significantly different (p < 0.05).

Considering that mTORC2 impairs proinflammatory signaling in macrophages^10^, we next investigated the impact of Rictor deletion in myeloid cells in liver disease induced by CDAHFD intake. As illustrated in Figure 5A-C, CDAHFD intake promoted body weight loss and reduced lean mass in both M-RicWT and KO mice, without impacting fat mass. Furthermore, CDAHFD intake similarly increased liver mass, promoted macrovesicular steatosis and augmented liver triacylglycerol and cholesterol contents in both M-RicWT and KO mice (Fig. 5D-G). Likewise, CDAHFD intake significantly reduced liver glycogen and protein contents, upregulated serum aminotransferases AST and ALT and diminished serum glucose, triacylglycerol, and cholesterol levels, regardless of mice genotype (Fig. 5H–N).

**Figure 5.**
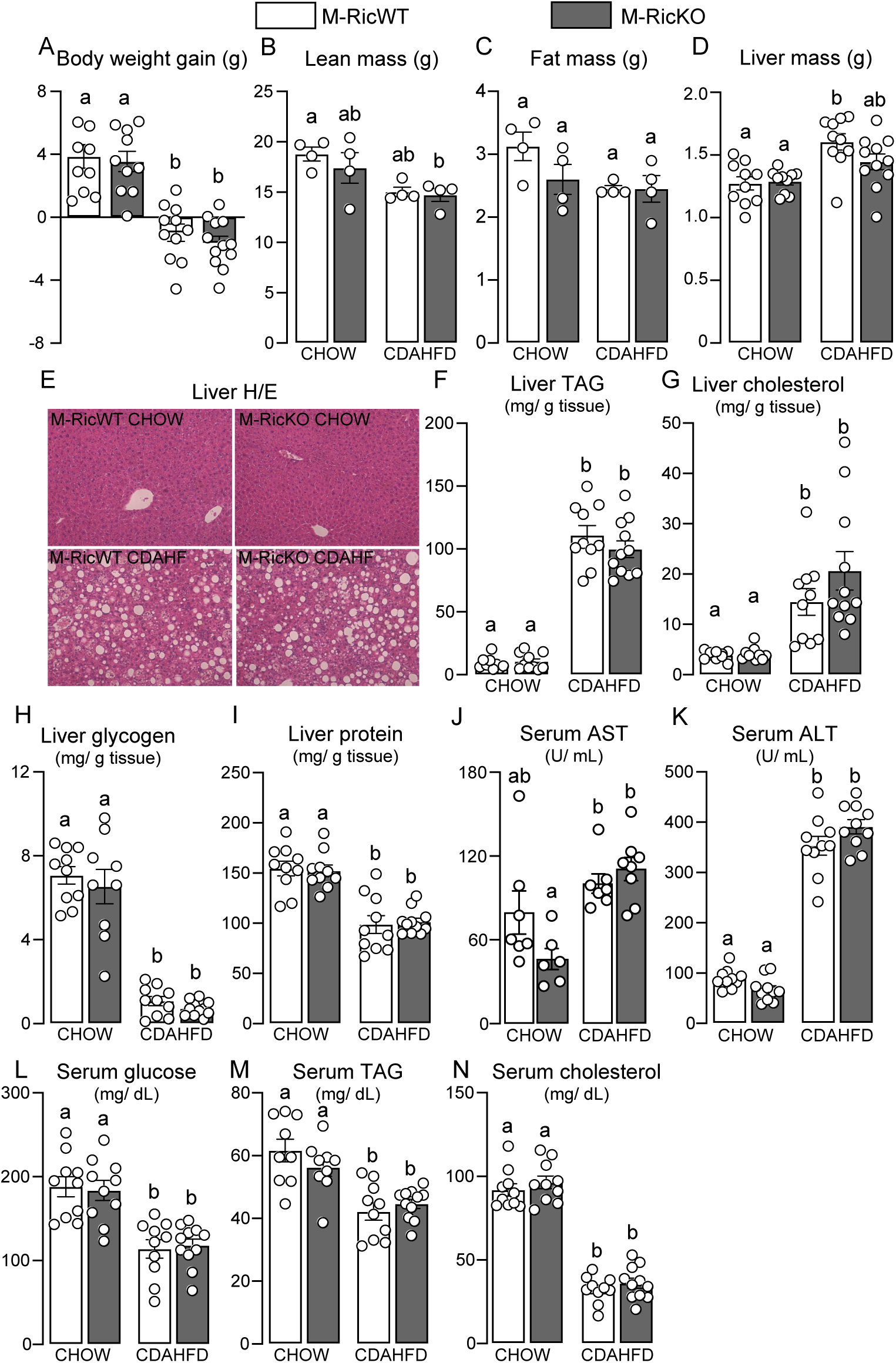
Myeloid cells mTORC2 deficiency has no impact on liver mass and serum metabolites in mice fed a CDAHFD for 10 weeks. Body weight gain (A), lean mass (B), fat mass (C), liver mass (D), representative H&E-stained liver sections (E), hepatic triacylglycerol (TAG, F), total cholesterol (TC, G), glycogen (H), and protein (I) contents, as well as serum aspartate aminotransferase (AST, J), alanine aminotransferase (ALT, K), glucose (L), TAG (M), and TC (N) levels in male Rictor flox (M-RicWT) and Rictor flox lysozyme M Cre^+^ (M-RicKO) mice fed either chow or choline-deficient, L-amino acid-defined, high-fat diet (CDAHFD) for 10 weeks. Data are presented as mean ± SEM. Two-way ANOVA followed by Tukey’s multiple-comparisons test. Groups that do not share a common superscript letter are significantly different (p < 0.05).

As previously observed in L-RicWT and KO mice (Fig. 3), CDAHFD intake increased hepatic IL-1β mRNA and protein contents and reduced liver TNF-α, IL-6, and IL-10 in similar magnitude in both M-RicWT and KO mice (Fig. 6C–E). Furthermore, CDAHFD intake increased liver mRNA content of macrophage markers *Adgre1* (F4/80) and *Cd86*, as well as the fibrosis-related genes *Col1a1*, *Tgfb1*, *Pdgfrb*, and *Timp1* independently of mice genotype, without significantly affecting mRNA levels of *Clec4f* and *Mrc1* (Fig. 6F-M).

**Figure 6.**
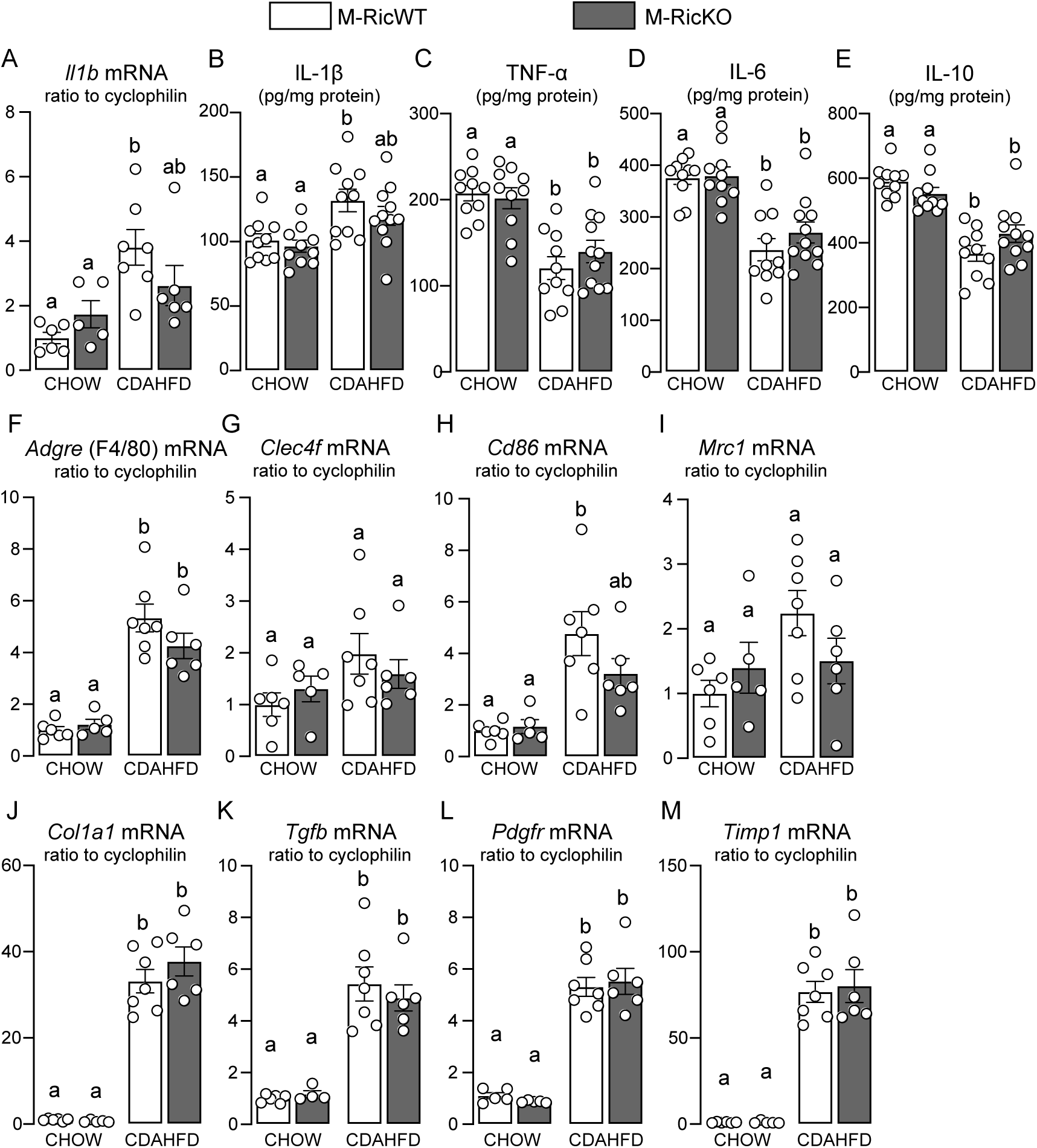
Myeloid cells mTORC2 deficiency does not affect liver inflammatory profile and pro-fibrotic gene expression in mice fed a CDAHFD for 10 weeks. Hepatic interleukin 1β (*Il1b*) gene expression (A) and protein content (B), tumor necrosis factor α (TNFα, C), IL-6 (D) and IL-10 (E) protein content and hepatic gene expression of *Adgre* F4/80 (F), *Clec4f* (G), *Cd86* (H), mannose Receptor C-Type 1 (*Mcr1*, I), collagen 1a1 (*Col1a1*, J), transforming growth factor β (*Tgfb*, K), *platelet-derived growth factor receptor* β (*Pdgfrb*, L) and tissue inhibitor of metalloproteinases (*Timp1*, M) in male Rictor flox (M-RicWT) and Rictor flox lysozyme M Cre^+^ (M-RicKO) mice fed either chow or choline-deficient, L-amino acid-defined, high-fat diet (CDAHFD) for 10 weeks. Data are presented as mean ± SEM. Two-way ANOVA followed by Tukey’s multiple-comparisons test. Groups that do not share a common superscript letter are significantly different (p < 0.05).

## Discussion

We investigated herein whether mTORC2 deficiency in distinct hepatic cell populations, namely hepatocytes and myeloid cells, including Kupffer cells and recruited macrophages, influences the development of liver disease induced by intake of a choline-deficient, amino acid-defined high-fat diet (CDAHFD), a mouse model in which liver disease is induced by impaired hepatic secretion of very low-density lipoprotein (VLDL) triacylglycerol. Our main findings indicate that hepatocyte Rictor/mTORC2 deficiency slightly attenuated CDAHFD-induced increases in liver mass, macrovesicular steatosis and triacylglycerol accumulation, without affecting though liver cholesterol, serum markers of liver injury (AST and ALT), as well as the upregulation in proinflammatory cytokine IL-1β and expression of fibrosis-related genes. Myeloid cells-Rictor deletion, which includes resident Kupffer cells and recruited macrophages, had no detectable impact on liver steatosis, inflammation, or fibrosis induced by CDAHFD. Altogether these findings indicate that hepatocyte mTORC2 deficiency has only slight, negligible role protecting mice from liver disease induced by CDAHFD intake.

Hepatic mTORC2 has been shown, via Akt phosphorylation at serine 473, to play a central role in promoting liver DNL through several different mechanisms including the upregulation of the transcriptional factor SREBP1c, which promotes the expression of DNL-related enzymes^8^. As expected from this prominent role in the regulation of DNL, hepatocyte-specific Rictor/mTORC2 deficiency completely abolished liver fat accumulation and disease induced by constitutive activation of PI3K-Akt signaling and DNL due to Pten deletion in hepatocytes^22^. Less striking protective effects of hepatocyte mTORC2 deficiency were seen in a mouse model in which liver disease is induced by intake of high-fat diet^22^. In contrast, our findings indicate that when hepatic steatosis develops through mechanisms that are not primarily driven by DNL, hepatocyte mTORC2 deficiency has a negligible impact, slightly reducing liver mass and triacylglycerol accumulation but not affecting other features of liver disease including inflammation, injury and fibrosis.

Interestingly, considering that mTORC2 deficiency and CDAHFD intake independently reduced liver DNL without additive effects of their combination, the reduction in liver triacylglycerol content seen in CDAHFD-fed L-RicKO probably occurs through additional mechanisms beyond DNL. To explore this possibility, we investigated alternative pathways that could contribute to the reduction in hepatic triacylglycerol content, including oxidative metabolism, mitochondrial mass and content of OXPHOS components. None of these parameters, however, were significantly altered in CDAHFD-fed L-RicKO, indicating that other mTORC2-dependent mechanisms, which will be addressed in future experiments, may underlie the observed reduction in hepatic lipid accumulation.

In addition to the regulation of lipid metabolism, previous studies have demonstrated that mTORC2 acts as a key negative regulator of proinflammatory signaling in macrophages^10^. Based on these findings, we investigated whether the deletion of Rictor in myeloid cells could influence the development of liver disease. Our results, however, showed that the absence of Rictor in myeloid cells did not affect the development of CDAHFD-induced liver disease in any of the parameters analyzed.

Inflammation plays a central role in the development and progression of MASLD, as inflammatory signals promote the activation of hepatic stellate cells, which are the main source of collagen deposition and fibrosis in the liver^23^. Although the CDAHFD model is widely recognized in literature as a model of MASH, we only found an increase in liver content of the proinflammatory cytokine IL-1β, which is produced by the NLRP3-inflammasome in response to several insults. Interestingly, CDAHFD intake reduced liver content of the classical proinflammatory cytokines TNF-α and IL-6, as well as the anti-inflammatory cytokine IL-10. Notably, the study that originally characterized this model did not assess liver content of these cytokines. Furthermore, histological inflammation scores increased during the early stages of disease development, but progressively declined with prolonged dietary exposure^12^. These findings suggest that the inflammatory response in the CDAHFD model is dynamic and stage-dependent, potentially reflecting a transition from an active inflammatory phase to a more advanced fibrotic stage, which may explain the reduced expression of both pro- and anti-inflammatory cytokines observed at the later time points.

In conclusion, hepatocyte mTORC2 deficiency show only modest effects in counteracting liver disease induced by CDAHFD intake only slightly reducing liver triacylglycerol content. Further studies are required to investigate the mechanisms underlying these mTORC2 actions that do not seem to depend on the regulation of liver DNL.

## Acknowledgements

This work was supported by grants from Sao Paulo Research Foundation (FAPESP #15/19530-5, 19/01763-4, 20/04159-8, 22/11234-1, 25/04262-7) and Brazilian National Council for Scientific and Technological Development (CNPq, #303784/2022-9) to WTF. Bianca F. Leonardi (#2021/14419-0), Ana B. Pires (#24/09406-4), Marina A. Abe-Honda (#24/13040-5), Álbert S. Peixoto (#22/02123-1), Érique Castro (#26/01093-2), Thayna S. Vieira (#23/04753-5), Natália M. Pessoa (#23/04509-7), Erika V. M. Pessoa (#24/12973-8), Ana Carolina P. Baptista (#23/17140-1), Loreana Silveira (#24/11195-1), and Mariana Mesquita (#24/16241-1) were recipients of fellowships from FAPESP. Natália Pontara-Corte CAPES was recipient of fellowship from Coordenação de Aperfeiçoamento de Pessoal de Nível Superior (CAPES).

## References

1. Tacke, F. et al. EASL–EASD–EASO Clinical Practice Guidelines on the management of metabolic dysfunction-associated steatotic liver disease (MASLD). J. Hepatol. 81, 492–542 (2024).

2. Younossi, Z. M. et al. The global epidemiology of nonalcoholic fatty liver disease (NAFLD) and nonalcoholic steatohepatitis (NASH): a systematic review. Hepatology 77, 1335–1347 (2023).

3. Sato-Espinoza, K., Chotiprasidhi, P., Liza, E., Placido-Damian, Z. & Diaz-Ferrer, J. Evolution of liver transplantation in the metabolic dysfunction-associated steatotic liver disease era: Tracking impact through time. World Journal of Transplantation vol. 14 Preprint at 10.5500/wjt.v14.i4.98718 (2024).

4. Harrison, S. A. et al. A Phase 3, Randomized, Controlled Trial of Resmetirom in NASH with Liver Fibrosis. New England Journal of Medicine 390, 497–509 (2024).

5. Sanyal, A. J. et al. Phase 3 Trial of Semaglutide in Metabolic Dysfunction–Associated Steatohepatitis. New England Journal of Medicine 392, 2089–2099 (2025).

6. Bansal, S. K. & Bansal, M. B. Pathogenesis of MASLD and MASH – role of insulin resistance and lipotoxicity. Aliment. Pharmacol. Ther. 59, S10–S22 (2024).

7. Targher, G., Byrne, C. D. & Tilg, H. MASLD: a systemic metabolic disorder with cardiovascular and malignant complications. Gut vol. 73 691–702 Preprint at 10.1136/gutjnl-2023-330595 (2024).

8. Hagiwara, A. et al. Hepatic mTORC2 activates glycolysis and lipogenesis through Akt, glucokinase, and SREBP1c. Cell Metab. 15, 725–738 (2012).

9. Laplante, M. & Sabatini, D. M. MTOR signaling in growth control and disease. Cell vol. 149 274–293 Preprint at 10.1016/j.cell.2012.03.017 (2012).

10. Festuccia, W. T., Pouliot, P., Bakan, I., Sabatini, D. M. & Laplante, M. Myeloid-specific rictor deletion induces M1 macrophage polarization and potentiates in vivo pro-inflammatory response to lipopolysaccharide. PLoS One 9, (2014).

11. De Ponti, F. F., Liu, Z. & Scott, C. L. Understanding the complex macrophage landscape in MASLD. JHEP Reports vol. 6 Preprint at 10.1016/j.jhepr.2024.101196 (2024).

12. Matsumoto, M. et al. An improved mouse model that rapidly develops fibrosis in non-alcoholic steatohepatitis. Int. J. Exp. Pathol. 94, 93–103 (2013).

13. Van Handel, E. Microseparation of Glycogen, Sugars, and Lipids. ANALYTICAL BIOCHEMISTRY vol. 11 (1965).

14. Festuccia, W. T. et al. The PPARγ agonist rosiglitazone enhances rat brown adipose tissue lipogenesis from glucose without altering glucose uptake. Am. J. Physiol. Regul. Integr. Comp. Physiol. 296, (2009).

15. Chaves-Filho, A. B. et al. Futile cycle of β-oxidation and de novo lipogenesis are associated with essential fatty acids depletion in lipoatrophy. Biochim. Biophys. Acta Mol. Cell Biol. Lipids 1868, (2023).

16. Castro, É. et al. Adipocyte-specific mTORC2 deficiency impairs BAT and iWAT thermogenic capacity without affecting glucose uptake and energy expenditure in cold-acclimated mice. Am. J. Physiol. Endocrinol. Metab. 321, E592–E605 (2021).

17. Bradford, M. M. A Rapid and Sensitive Method for the Quantitation of Microgram Quantities of Protein Utilizing the Principle of Protein-Dye Binding. ANALYTICAL BIOCHEMISTRY vol. 72 (1976).

18. Chimin, P. et al. Adipocyte mTORC1 deficiency promotes adipose tissue inflammation and NLRP3 inflammasome activation via oxidative stress and de novo ceramide synthesis. J. Lipid Res. 58, 1797–1807 (2017).

19. Guillot, A. et al. Mapping the hepatic immune landscape identifies monocytic macrophages as key drivers of steatohepatitis and cholangiopathy progression. Hepatology 78, 150–166 (2023).

20. Guillot, A., Kohlhepp, M. S., Bruneau, A., Heymann, F. & Tacke, F. Deciphering the immune microenvironment on a single archival formalin-fixed paraffin-embedded tissue section by an immediately implementable multiplex fluorescence immunostaining protocol. Cancers (Basel). 12, 1–18 (2020).

21. Borges, M. C. et al. Obesity-induced hypoadiponectinaemia: The opposite influences of central and peripheral fat compartments. Int. J. Epidemiol. 46, 2044–2055 (2017).

22. Yuan, M., Pino, E., Wu, L., Kacergis, M. & Soukas, A. A. Identification of Akt-independent regulation of hepatic lipogenesis by mammalian target of rapamycin (mTOR) complex 2. Journal of Biological Chemistry 287, 29579–29588 (2012).

23. Hammerich, L. & Tacke, F. Hepatic inflammatory responses in liver fibrosis. Nature Reviews Gastroenterology and Hepatology vol. 20 633–646 Preprint at 10.1038/s41575-023-00807-x (2023).

